# Directed, but not random, breast cancer cell migration is faster in the G1 phase of the cell cycle in 2D and 3D environments

**DOI:** 10.1101/288183

**Authors:** Kamyar Esmaeili Pourfarhangi, Edgar Cardenas de la Hoz, Andrew R. Cohen, Bojana Gligorijevic

**Affiliations:** Bioengineering department, College of Engineering, Temple University, Philadelphia, PA, 19122; Department of Electrical and Computer Engineering, College of Engineering, Drexel University, Philadelphia, PA, 19104; Cancer Biology Program, Fox Chase Cancer Center, Philadelphia, PA, 19111

## Abstract

Cancer cell migration is essential for the early steps of metastasis, during which cancer cells move through the primary tumor and reach the blood vessels. In vivo, cancer cells are exposed to directional guidance cues, either soluble, such as gradients of growth factors, or insoluble, such as collagen fiber alignment. Depending on the number and strength of such cues, cells will migrate in a random or directed manner. Interestingly, similar cues also stimulate cell proliferation. In this regard, it is not clear whether cell cycle progression affects migration of cancer cells and whether this effect is different in random versus directed migration. In this study, we tested the effect of cell cycle progression on random and directed migration, both in 2D and 3D environments, in the breast carcinoma cell line, FUCCI-MDA-MB-231, using computational image analysis by LEVER. Directed migration in 2D was modeled as chemotaxis along a gradient of soluble EGF inside 10 µm-wide microchannels. In 3D, directed migration was modeled as contact guidance (alignotaxis) along aligned collagen fibers. Time-lapse recordings of cells in 2D and 3D revealed that directed, but not random migration, is cell cycle-dependent. In both 2D and 3D directed migration, cells in the G1 phase of the cell cycle outperformed cells in the G2 phase in terms of migration persistence and instantaneous velocity. These data suggest that in the presence of guidance cues *in vivo*, breast carcinoma cells in the G1 phase of the cell cycle may be more efficient in reaching vasculature.

## Introduction

Cancer is one of the leading causes of deaths globally. Cancer mortality is tightly linked to invasion and metastasis, for which no treatment exists at the moment.^1^ During metastasis, tumor cells invade and migrate through the stromal tissue, disseminate via the lymphatic or vascular systems and colonize distant organs.^2^ While migrating through the tissue, tumor cells are exposed to guidance cues, resulting in directed migration that facilitates persistent navigation through the tissue and efficient arrival to the lymphatic or blood vessels.^3^ Directed migration is guided by biochemical and biophysical cues. The best studied soluble cues are gradients of growth factors and cytokines which induce chemotaxis.^4^ Recent studies of extracellular matrix (ECM) properties have identified a number of biophysical cues, one example of which is alignment of collagen fibers, shown to stimulate contact guidance i.e. alignotaxis.^5,6^

In the case of mesenchymal cancer cells, both random and directed migration are composed of four interdependent molecular steps that make the cell motility cycle.^7^ The first step involves formation of adhesive protrusions at the leading edge, whose direction is determined by cell polarization.^3^ The next steps include formation of new adhesions at the cell front, elevation of actomyosin contractility in the cell body, and retraction of the adhesions at the cell rear. Cues that induce directed migration, such as growth factor gradients or aligned collagen fibers, cause local activation of certain Rho family GTPases that are part of the polarity signaling machinery of the cells.^6^ Hence, cells are persistently polarized in the direction of the guiding cues, and therefore, new adhesive protrusions are formed in the same direction resulting in a guided migration.^8,9^ A forthright consequence of directional migration is an increase in persistence, which is also manifested in higher average speed of directionally migrating cells.

Based on the established hallmarks of cancer,^10^ it can be expected that motile cancer cells are actively cycling and proliferating. However, it is unlikely that cells actively engaged in actin reorganization during migration can proceed through cell cycle and division due to both structural and energy constraints. In support of this thinking, a number of studies has reported that tumors which invade and metastasize more efficiently, grow slower and vice versa.^11–13^. In melanoma cancers, this phenomenon was described as the *phenotypic switch*^11^. To explain how the switch between proliferation and invasion may occur at a single cell level, the Go-or-Grow hypothesis was proposed, suggesting a temporal exclusivity in division and motility.^14^ However, testing of this hypothesis was mainly done as ensemble measurements and so far, resulted in contradicting reports.^15–17^ For example, in glioma and cervical cancer cell lines, cells in G1 and S phase demonstrate higher speed of 2D random migration.^18^ Next, melanoma cells in 3D spheroids show little migration and invasion in G1 phase compared to S/G2, while colon cancer cells *in vivo* were shown to migrate faster when in S/G2/M phase compared to G1.^19^ While the current literature suggests that the cell cycle phase may influence cell speed and extent of cell invasion, a systematic study of random and directed cell migration in 2D and 3D in breast carcinoma cells was not done.

Although cancer cell migration in vivo is mostly directional and guided by soluble and physical cues, the effect of cell cycle on directed migration was not previously tested. In this study, we tested the cell cycle-dependency of persistence and velocity in random and directed migration, in 2D and 3D models. To monitor the cell cycle status real-time, we utilized the nuclear labeling by FUCCI fluorescent set,^20^ which allows us to distinguish the G1 from the S/G2 phase of the cell cycle. Using LEVER for computational image analysis, we were able to automate segmentation, tracking and analysis of migrating cells. Our results indicate that in both 2D and 3D conditions, only directed migration is affected by the cell cycle phase. While no cell cycle dependency was observed for either persistence or velocity in randomly migrating cells, directionally migrating cells in the G1 phase of the cell cycle were faster (in 2D and 3D) and more persistent (3D) than cells in the S/G2 phase of the cell cycle. Taken together, our results suggest that cells in the G1 phase of the cell cycle excel at directed migration.

## Results

### Establishment and characterization of a 2D model for directed cell migration

To model directed cell migration in 2D, we designed a microchip with microchannels with a gradient of EGF as chemoattractant (Figure 1A).^21,22^ Briefly, two reservoirs were created by a biopsy punch on each side of a 20-microchannel array (W 10 µm × H 20 µm × L 850µm). The left reservoir was loaded with cells (Figure 1A) and the right reservoir acted as the chemoattractant source. To facilitate cell entry into the channels, the left end of the microchannels was tapered (W 35 µm × L 150µm). The 10 µm-wide microchannels guide migration of MDA-MB-231 cells along x-axis, without cell constriction.^23^

**Figure 1.**
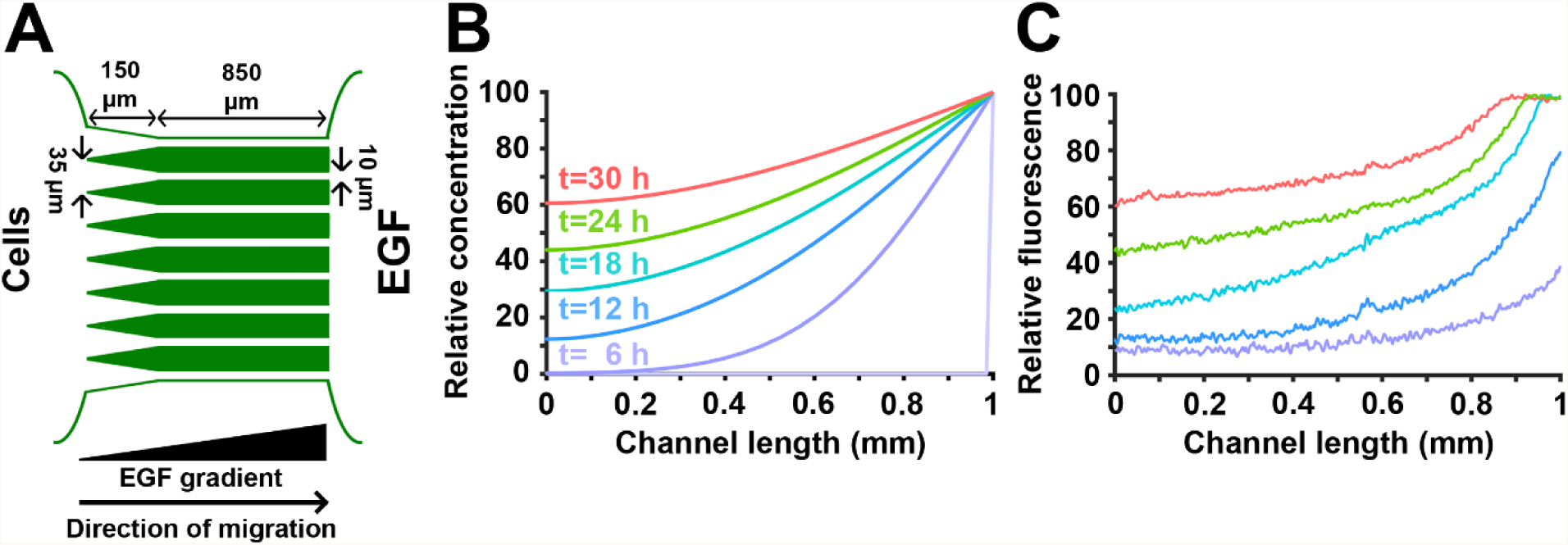
Development and characterization of a 2D model for directed cancer cell migration using a chemoattractant gradient in microchannels. **A.** Simplified microchip design. To facilitate cell entry into 10 µm-wide microchannels, left end of microchannels was tapered (35 µm width). **B.** Mathematical modeling of the non-linear chemoattractant gradient across the microchannels. Example time points ranging from 6-30h are shown. **C.** Gradient of AlexaFluor 405 dextran across the microchannel length, measured at different times 6-30h. Colors correspond to acquisition times indicated in (**B**).

To test the microchip ability to provide a stable gradient across its full length, formation and dynamics of the gradient was first mathematically simulated by solving the unsteady-state diffusion equation using a finite element approach. **Movie S1** and Figure 1B show the results of the mathematical modeling and formation of the gradient across the microchannels. As indicated in Figure 1B, the gradient steepness is predicted to be stable from 12 to 18 h post-gradient formation.

We next experimentally validated the mathematical model by monitoring diffusion of fluorescently labeled dextran over time. AlexaFluor 405-labeled 10 kDa dextran has a molecular weight and diffusivity similar to those of EGF (MW = 6 kDa and D = 0.5e-4 µm^2^/s). Mass transfer was limited to molecular diffusion by capping reservoirs with PDMS pieces, equilibrating the pressure across the system and reducing convection. Figure 1C shows quantification of gradient profiles at different time points. While there is a reduction in the gradient steepness over time, as mathematically shown, average gradient steepness in the system is stable from 12 to 18 h post-gradient formation (< 10% difference). Since the chemotaxis of MDA-MB-231 cells is affected by the steepness and concentration of the EGF,^24^ quantification of the cell migration parameters (instantaneous velocity and migration persistence) was only conducted on data acquired from 12 to 18 h post-gradient formation.

### Directed, but not random, migration of cancer cells in 2D is cell cycle-dependent

We first tested the cell cycle-dependency of cancer cell migration in 2D by acquiring time-lapse recordings of FUCCI-MDA-MB-231 cells (**Movie S2**). To model 2D random migration, cells were plated on gelatin-coated dishes. In the directed cell migration model, cells moved following the chemoattractant onto the gelatin-coated microchannels and allowed to migrate along the chemoattractant gradient (**Movie S3**).

Cell trajectories and the corresponding cell cycle phase information were extracted from time-lapse movies via LEVER (see Materials and methods) (**Movie S4)** and used to calculate cell persistence and instantaneous velocity.

Figure 2A and 2B depict representative trajectories of cells randomly migrating in 2D and the corresponding cell cycle phases (G1, red and S/G2, green). Cells in the early S phase of the cell cycle express both red and green fluorophores, generating yellow labeling. In our system, these cells amounted to <10% of the total number of cells and were excluded from the quantification. As expected, without guiding cues, cells freely change their polarization direction, resulting in a very low migration persistence, independently from the cell cycle phase (Figure 2C).^3^ Consequently, instantaneous velocity of cells in the G1 phase of the cell cycle shows no significant difference compared to that of cells in the S/G2 phase of the cell cycle (Figure 2D). Taken together, **these findings** demonstrate that random migration of cancer cells in 2D is not cell cycle-dependent.

**Figure 2.**
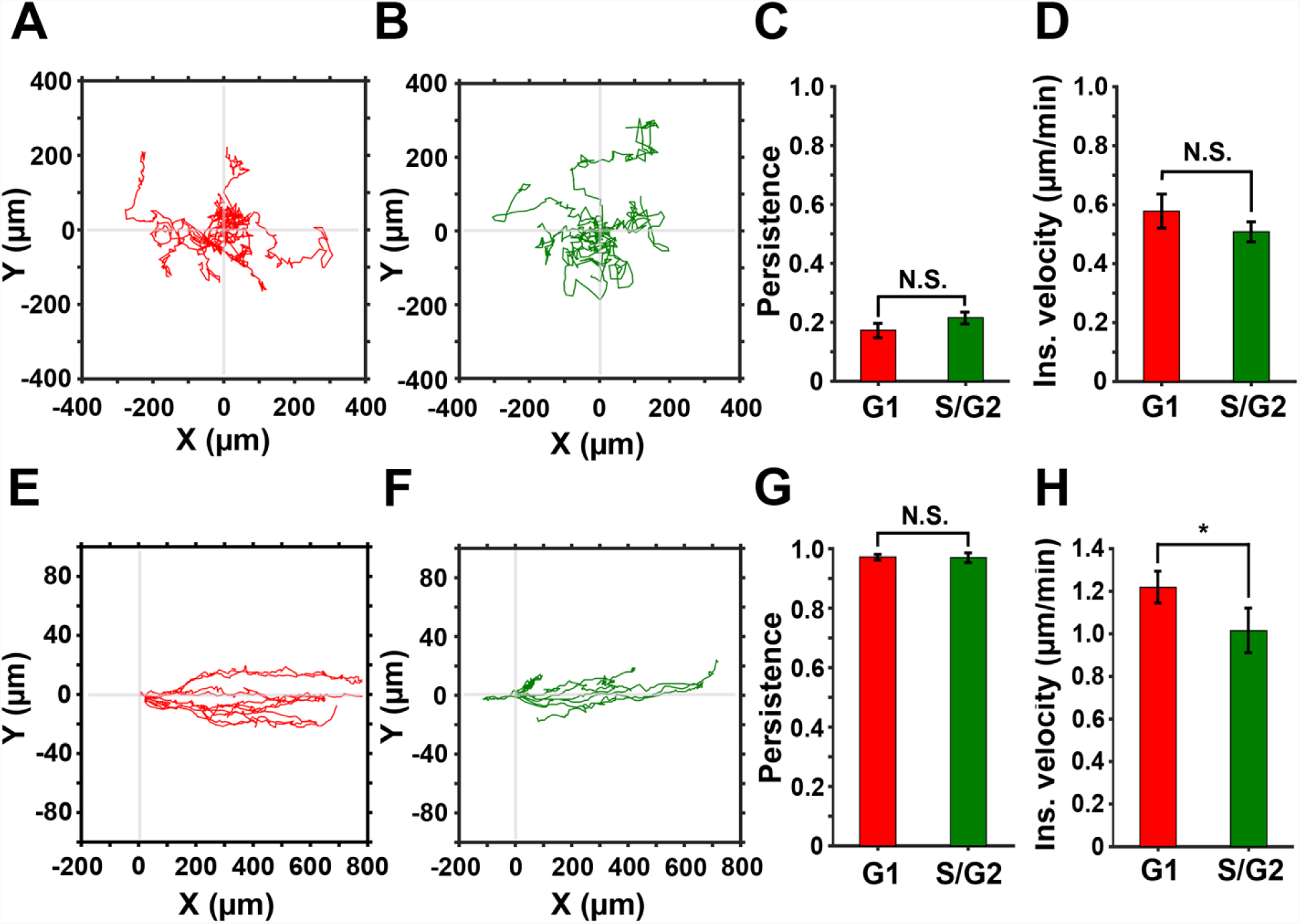
2D directed, but not random, migration is cell cycle-dependent. **A & B.** Representative trajectories of cells in the G1 (red) and the S/G2 (green) phase of the cell cycle, randomly migrating on gelatin-coated plates. **C & D.** Persistence and instantaneous velocity of cells randomly migrating in 2D. **E & F.** Representative trajectories of cells in the G1 (red) and the S/G2 (green) phases of the cell cycle directionally migrating inside the gelatin-coated microchannels. **G & H.** Persistence and instantaneous velocity of cells directionally migrating inside microchannels.

Next, we assessed cells subjected to directional migration in 2D (Figure 2E-H). Microchannels guide the cell migration along x-axis without applying spatial constriction,^23^ while the non-linear gradient of EGF stimulates chemotaxis in MDA-MB-231 cells,^24^ ensuring unidirectional movement. Simultaneous exposure to physical confinement and gradient of chemoattractant results in highly persistent cell migration, similar in both the G1 and the S/G2 phase of the cell cycle (Figure 2G). However, instantaneous velocity of cells in G1 is significantly higher (approximately 20%) than the velocity of cells in S/G2 phase of the cell cycle (Figure 2H). Collectively, our data suggest that directed migration of cancer cells in 2D is cell cycle-dependent. While migration persistence of cells in the G1 and S/G2 phases is comparable, cells in the G1 phase exhibit significantly higher velocity.

### Directed, but not random, migration of cancer cells in 3D is cell cycle-dependent

A number of studies demonstrated that cell migration in 2D may not reflect cell behaviors in more physiological relevant 3D models, where cells can be exposed to tissue-like confinement^25–27^ and reciprocal cell-matrix interactions^28^ One such model is the acid-soluble collagen gel, which provides fibrillar structure resembling the topography of tissue ECM, and therefore, is superior to the homogeneous 3D gels.^29^ In 3D collagen gels with pore size equal to or larger than the diameter of cell nucleus, breast cancer cells utilize MMP-independent migration. In contrast, MMP-dependent invasion is present in environments with smaller pores.^30^ Additionally, directed cell migration along aligned and bundled collagen fibers (contact guidance) was shown to significantly increase persistence,^5^ without affecting the instantaneous velocity of cells.^31^

To test the cell cycle-dependency of random and directed cell migration in 3D, we generated collagen gels with either randomly oriented (Figure 3A-C) or aligned fibers (Figure 3D-F). The aligned fiber architecture was achieved by flowing magnetic beads through the collagen gel.^32^ We confirmed the orientation of the fibers by confocal reflection imaging (Figure 3B, 3E) and measured the distribution of fiber angles in each condition. The randomly oriented fibers show a uniform distribution of fiber angles (Figure 3C), indicating that there is no enrichment of fibers in any particular angle. The aligned fibers show a Gaussian distribution of angles, indicating that the majority of fibers are positioned at the same angle (Figure 3F).

**Figure 3.**
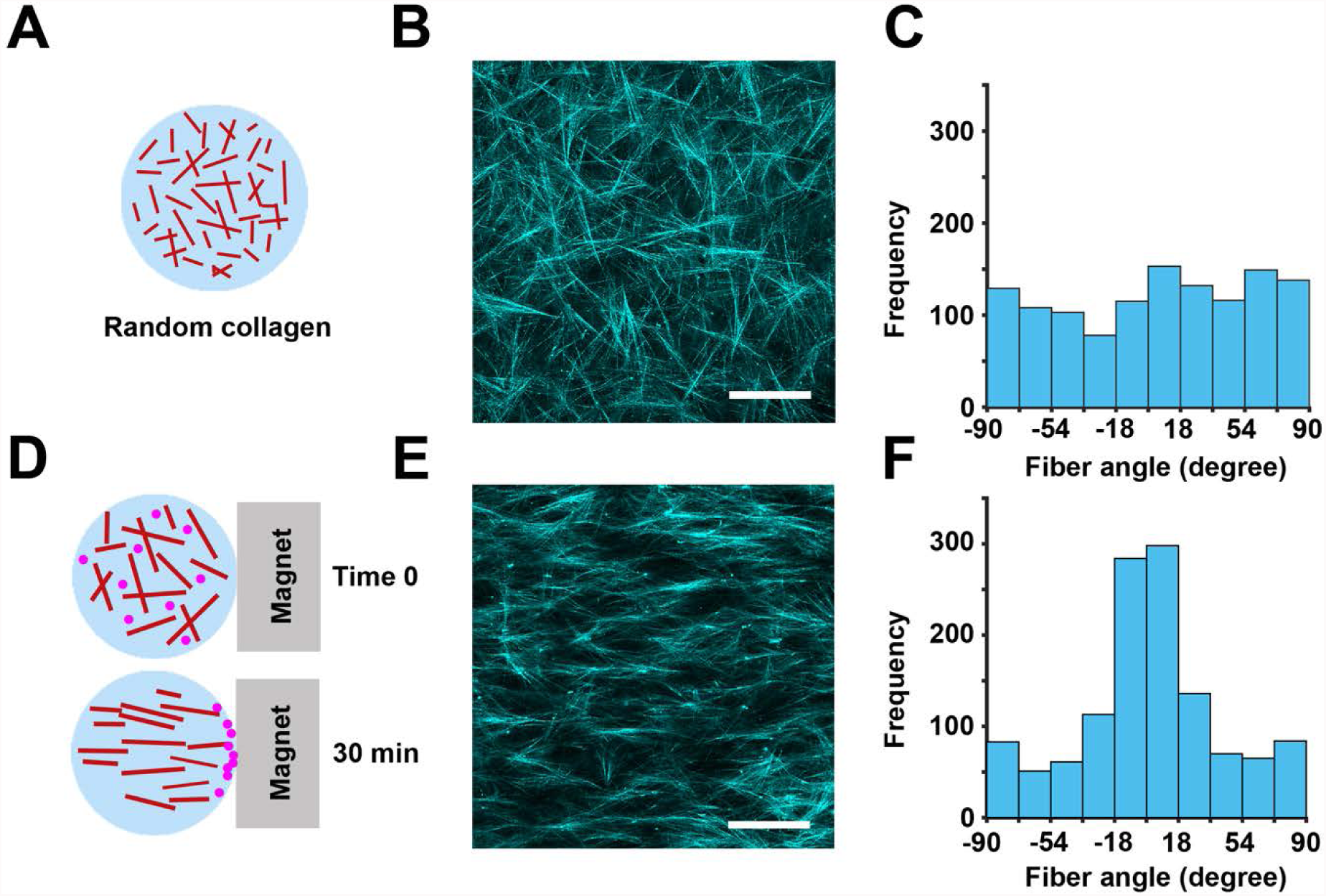
Development of a 3D model for directed cell migration using collagen fiber alignment. **A.** Randomly oriented 3D fibrillar collagen was used in the 3D random migration assay. **B.** Representative image of the random architecture of collagen fibers imaged by confocal reflection microscopy. **C.** Quantification of collagen fiber angles in (**B**) demonstrating a random distribution of the collagen fiber orientation. **D.** Collagen mixed with magnetic beads and exposed to magnetic-induced flow results in 3D collagen with aligned fibers. **E.** Aligned architecture of collagen fibers imaged by confocal reflection microscopy. **F.** Distribution of collagen fiber angles in (**E**) shows a Gaussian distribution in collagen fibers direction, centered at 90° relative to the magnet orientation. Scale bar 100 µm.

Using these 3D models, we compared the migration of FUCCI-MDA-MB-231 cells in randomly oriented (**Movie S5**) or aligned (**Movie S6**) fibrillar collagen. Time-lapse images were segmented and tracked using LEVER. Consistent with the random migration pattern in 2D, cell cycle status did not affect neither persistence nor velocity of cells in randomly oriented fibers in 3D (Figure 4A-D).

**Figure 4.**
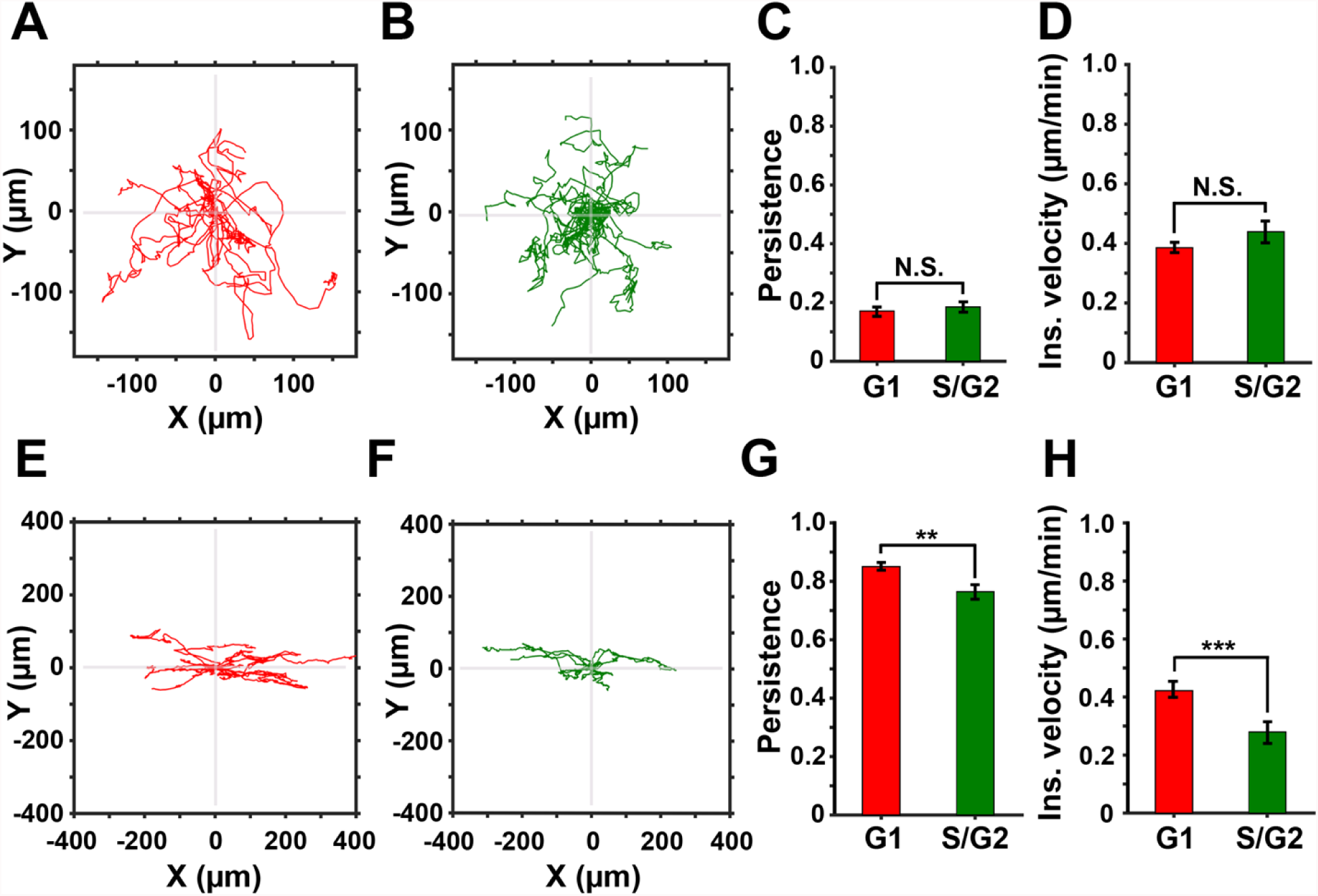
3D directed, but not random, migration is cell cycle-dependent. **A & B.** Representative trajectories of cells in the G1 (red) and the S/G2 (green) phase of the cell cycle migrating in 3D collagen with randomly oriented fibers. **C & D.** Persistence and instantaneous velocity of cells migrating in 3D collagen with randomly oriented fibers. **E & F.** Representative trajectories of cells in the G1 (red) and the S/G2 (green) phase of the cell cycle migrating in aligned 3D collagen fibers. **G & H.** Persistence and instantaneous velocity of cells directionally migrating in aligned 3D collagen fibers.

In aligned collagen, the trajectories show higher persistence compared to cells in randomly oriented collagen (Figure 4E and 4F). Quantification of the migration parameters shows that cells in the G1 phase of the cell cycle are both significantly faster and more persistent compared to the cells in S/G2 of the cell cycle (Figure 4G and H). In conclusion, our data demonstrate that cells in the G1 phase of the cell cycle exhibit faster migration in response to guidance cues, i.e. directed migration. During random migration, persistence and velocity of cells in the G1 and S/G2 phases of the cell cycle are similar.

## Discussion

In this study, we sought to determine the migration parameters of cancer cells relative to their cell cycle status. To this purpose, we established models of 2D and 3D directed migration and utilized MDA-MB-231 breast carcinoma cells expressing the FUCCI cell cycle reporter. Using time-lapse live imaging, we demonstrate that only the directed motility of cancer cells changes throughout the cell cycle progression. We showed that under directed guidance cues cells in G1 outperform those in S/G2 by exhibiting higher persistence and velocity both in 2D and 3D environments.

The persistence and the velocity of cells can be regulated by extrinsic (ECM properties, chemoattractant gradient, interstitial flow) ^24,33–35^ as well as intrinsic cues (expression levels of receptor tyrosine kinases, integrins, metabolism). In the current study, we keep the external environment stable, which allows us to isolate the relationship between the intrinsic cell cycle signaling and migration.

The cross-talk between cell cycle and motility pathways was suggested by a number of mechanistic reports. For example, integrins and receptor tyrosine kinases were shown to regulate both RhoGTPases, master regulators of cell contractility and actin polymerization,^36^ as well as G1-related cyclin-dependent kinases.^37^

Our models and method provide a platform for further interrogating the relationship between the cell cycle progression and other extrinsic cues and the effects of the interplay not only on cell migration, but also on cell invasion. For instance, ECM stiffening in mammary epithelial cells activates cell cycle progression via a FAK-Rac-Cyclin D1 pathway,^38^ while applying sheer stress to tumor cells can induce a G2/M arrest through α_v_ß_3_ and ß_1_-mediated pathways.^39^ Furthermore, similar to cell migration, invasion and ECM degradation can be modulated by external factors, such as soluble cues and ECM parameters.^40–43^ In addition, intrinsic cellular activities, such as cell cycle, can also affect the ECM degradation. Recent study has shown that the cytoplasmic pool of cyclin-dependent kinase inhibitor p27 is involved in regulation of ECM degradation.^44^ On a similar note, anchor cell invasion into vulval epithelium, occurring during development in *C. elegans*, is only performed by cells that are in G0/G1 phase of the cell cycle.^45^

Tumor cell arrest during the G1/S transition, by inhibitors of Cdk4/6, is one of the new avenues for breast carcinoma treatment.^46^ While such treatment successfully reduces tumor size, it may also accumulate reactive oxygen species and increase Cyclin D1 expression,^47^ which have been linked to increased migration and invasion.^48,49^ Hence, understanding of the coordination between cell cycle and cell migration, as well as the other hallmarks of metastasis is increasingly important.

## Methods

### Cell culture and generation of FUCCI-MDA-MB-231

Human breast cancer cell line MDA-MB-231 (ATCC, Manassas, VA) was cultured in DMEM supplemented with 10% FBS (Atlanta Biologicals, Flowery Branch, GA) and 1% Penicillin/Streptomycin mixture (Gibco, Thermofisher Scientific, Waltham, MA).

Cell line FUCCI-MDA-MB-231^20^ was generated using a self-inactivating lentiviral expression vector system (Miyoshi j virol 1998).^50^ Viral particles were produced by co-transfecting mKO2-hCdt1 (red) or mAG-hGem (green) with the packaging plasmids (pCAG-HIVgp), G protein of vesicular stomatitis virus (VSV-G) and Rev-expressing plasmid (pRSV-rev) into HEK-293T cells. Supernatant was collected and concentrated using the Lenti-X concentrator (Clontech Laboratories, Inc). High-titer viral solutions were used to co-transduce MDA-MB-231 cells and top 5% expressors were selected by FACS.

### 2D migration assay

60,000 FUCCI-MDA-MB-231 cells were cultured on gelatin-coated plates described previously.^51^ Briefly, acid-washed 35-mm glass bottom dishes (MatTek Corporation, Ashlan, MA) were incubated with 50 µg/ml Poly-L-Lysin (Gibco, Thermofisher Scientific, Wlatham, MA) for 20 min and then, coated with 0.2% gelatin solution for 10 min. Plates were then washed with PBS (Gibco, Thermofisher Scientific, Waltham, MA) and cross-linked by 0.2% glutaraldehyde (GTA, Sigma, St. Louis, MO) on ice for 15 min. Next, plates were extensively rinsed with PBS, quenched with 5 mg/ml sodium borohydride (Sigma-Aldrich, St. Louis, MO) and sterilized with 70% ethanol (Decon Laboratories, King of Prussia, PA). Image acquisition was initiated 2 h after cell plating.

### Microchannels fabrication

The microchips containing microchannels were fabricated using soft lithography techniques. First, the microchip design was created in Autocad (Autodesk, San Rafael, CA) and printed on a glass plate to serve as a photomask. Next, a silicon master was fabricated via spin-coating a 4” silicon wafer (University Wafers, South Boston, MA) with SU-8 photoresist (MicroChem Corporation, Newton, MA) followed by baking at 70 °C for 20 min, exposing to UV light passing through the photomask transparencies and removing the uncross-linked photoresist by a developer. The silicon master served as a mold for making microchannels, onto which a 10:1 mixture of PDMS:curing agent (Dow Corning, Midland, MI) was poured, degassed under vacuum to remove air bubbles, and cured at 70 °C for 1 h. Next, cured PDMS was carefully peeled from the wafer and cut into single-chip size pieces. Cell- and chemoattractant-side reservoirs were made on each device using a 5-mm biopsy punch. Next, acid-washed 35-mm glass bottom dishes (MatTek Corporation, Ashland, MA) and PDMS devices were activated via oxygen plasma treatment for 30 seconds at 300 mTorr. Microchips were then assembled by bonding each PDMS device with a MatTek dish. Assembled devices were then sterilized with 70% ethanol followed by repeatedly rinsing the devices with sterile PBS.

### Mathematical modeling of gradient formation within the channels

Dynamics of EGF gradient formation within the microchannels were modeled in MATLAB (Mathworks, Natick, MA) using the partial differential equation toolbox (PDEtool). The geometry of the model included xy cross-section of a microchannel, and physics of the model involved a two-dimensional unsteady-state diffusion equation, 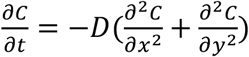, with a constant-concentration boundary condition at the source edge and a no-flux boundary condition applied to the sink edge of the channel as well as the edges in contact with PDMS. Initial concentration of the EGF in the whole system was set to zero. The EGF-water diffusion coefficient, D, was set to 0.5e-4 cm^2^/s.

### Experimental validation of gradient formation in the microchannels

Prior to the experiment, sink and source reservoirs of the devices were rinsed and filled with PBS. The PBS in the source reservoir was supplemented with Alexa Fluor 405-labeled 10 KDa dextran (MW of EGF is 6 kDa), and microchannels were then imaged on a widefield Olympus (Olympus, Tokyo, Japan) microscope for 30 hours with 10 min intervals.

Resulting movies of the dynamics of Alexa Fluor 405-dextran diffusion within the microchannels across the two reservoirs were analyzed in Fiji.^52^ briefly, a “plot profile” was applied to a line drawn through the length of the channels and fluorescence intensities were recorded at every frame. Relative fluorescence intensities were then calculated via applying the following equation to the raw data:

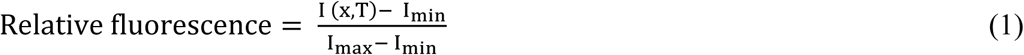

I(x,T) represents the raw fluorescence intensity recorded at length x of the microchannel and time T of the experiment. I_min_ and I_max_ are minimal and maximal fluorescence intensities recorded in the microchannel.

### 2D directed migration assay using microchannels

Prior to cell plating, sink and source reservoirs of the microchips were rinsed with PBS and then filled with culture media. 25,000 cells were plated in the cell (sink) reservoir and the devices were placed in incubator for 2 h for cell adhesion. Next, the media in the source reservoir was supplemented with 20 µM EGF (Thermofisher Scientific, Waltham, MA) and microchips were imaged with a widefield Olympus (Olympus, Tokyo, Japan) microscope for 30 hours with 10 min intervals.

### Collagen alignment

Collagen fibers were aligned by incorporating paramagnetic polystyrene beads (PM-20-10; Spherotech, Lake Forest, IL) into a 1.5 mg/ml collagen mixture at 4% (v/v) and exposing the mixture to the magnetic field of a neodymium magnet (BZX0Y0X0-N52; K&J Magnetic, Pipersville, PA) during collagen polymerization.^5,32^ This step was performed at room temperature for 30 min. As schematically shown in Figure 3D, magnetic-field induced flow of magnetic beads within the collagen matrix aligns collagen fibers. Collagen alignment was assessed by confocal reflection microscopy followed by image processing by ct-FIRE^53^ to extract fiber angles.

### 3D migration assays

40,000 FUCCI-MDA-MB-231 cells were suspended in 50 µl of a collagen mixture containing: 1.5 mg/ml rat tail collagen I (Corning, Tewksbury, MA), 5 µl 10X PBS, 10% FBS (Atlanta Biologicals, Flowery Branch, GA), 1% Penicillin/Streptomycin, DMEM (Gibco, Thermofisher Scientific, Waltham, MA), 4% paramagnetic polystyrene beads (PM-20-10; Spherotech, Lake Forest, IL) and 1 N NaOH. The mixture was vortexed at 4 °C for 5 min and pipetted in a 35-mm glass bottom plate (MatTek Corporation, Ashland, MA). Collagen mixture was polymerized at room temperature for 30 minutes. For collagen alignment, this step was conducted by positioning the plate adjacent to a neodymium magnet (BZX0Y0X0-N52; K&J Magnetic, Pipersville, PA). Next, 1ml DMEM containing 10% FBS and 1% antibiotics was pipetted into the plate. The plate was then imaged via a widefield Olympus IX81 (Olympus, Tokyo, Japan) microscope for 30 hours with 10 min intervals.

### Live cell imaging

Live cell imaging was performed via a widefield Olympus (Olympus, Tokyo, Japan) microscope equipped with LED lamp, Hamamatsu Orca 16-bit CCD (Hamamatsu, Hamamatsu city, Japan), automated z-drift compensation IX3-ZDC (Olympus, Tokyo, Japan), automated Prior stage (Prior Scientific, Rockland, MA) and an environmental chamber. For live cell imaging, regular culture media was supplemented with 1:100 Oxyfluor (Oxyrase, Mansfield, OH) and 10 mM sodium lactate (Sigma-Aldrich, St. Louis, MO) to reduce phototoxicity. Time lapse imaging was conducted at a single focal plane using an Olympus 20x 0.7 NA objective and images were acquired at FITC, TRITC, and bright field channels. Images were collected every 10 minutes for 30 hours.

### Computational image analysis

Each image is first segmented, and then tracked using the approach previously described for the LEVER (lineage editing and validation) software tools^54–57^. The segmentation processed the two FUCCI channels using a denoising algorithm^58^ that models imaging noise as a combination of slow-varying low frequency background noise and high frequency shot noise. The denoising algorithm uses a Gaussian low-pass filter, with kernel size equal to 10% of the image size, combined with a median filter with support of 3×3 pixels. The segmentation treats the fluorescence and phase channels separately. The fluorescence images start with an adaptive Otsu thresholding, followed by a connected component analysis. The phase segmentation identifies bright and dark foreground pixels as those falling greater than one standard deviation from the mid-level background pixels. These bright and dark pixels are combined with an Otsu thresholded gradient image to produce the final foreground. The intersection of the phase and fluorescence channel segmentations is used as the final segmentation.

Following segmentation, the images are tracked to establish temporal correspondences among the segmentation results. The multitemporal association tracking (MAT)^59,60^ algorithm has been widely applied in a number of applications and is used here. MAT uses a minimum spanning tree optimization to solve the data association problem across multiple image frames (here set to three) in polynomial time. The first tracking step is to compute a cost function between segmentations based on differences in spatial location, shape and size, and fluorescence intensity signals. For each cell, the fluorescent intensity is taken as the mean fluorescent signal within the segmentation boundaries. Cells that are separated by more than twice the maximum radius of either cell, or with a size difference of more than 90% between frames are gated or considered to have an infinite tracking cost. The gate is adaptive, with the constraint adjusted upwards until at least five possible tracking matches are obtained. MAT then uses the cost function to compute optimal tracking associations between segmentations. The LEVER program then allows the results to be visualized, and optionally for any errors to be corrected manually.

Cell tracks were manually analyzed to correct for mis-segmented cells and assess instantaneous velocity (displacement between two frames divided by time) and cell persistence (net cell displacement over the course of experiment divided by the total displacement).

### Ethics Statement

Ethics approval was not required for this work.

### Statistical analysis

Two-tailed student T-test was performed for all statistical analysis. Statistical significance was defined as * P < 0.05; ** P < 0.01 and *** P < 0.001. Data are shown as means ± SEM.

## Author Contributions

KE and BG have designed and interpreted all experiments. KE, BG, ECH and ARC analyzed the data and wrote the manuscript. KE has performed all experiments.

## Acknowledgements

We would like to thank Dr. Battuya Bayarmagnai and Mr. Louisiane Perrin for help in editing the manuscript. Authors declare no competing interests. This work was funded by NIH 5K99CA172360 and Concern Foundation grant to BG and NIH NINDS R01NS076709 and NIH NIA R01AG041861 to ARC.

