## Supplementary Materials for "Directed, but not random, breast cancer cell migration is faster in the G1 phase of the cell cycle in 2D and 3D environments"

### Supplementary Movies

**Movie S1. Mathematical simulation of EGF diffusion within the microchannels.**

The results of the mathematical simulation of the formation and dynamics of EGF non-linear gradient within the microchannels is shown. Colors correspond to the EGF concentration indicated in the legend. Time between frames is 5 hours.

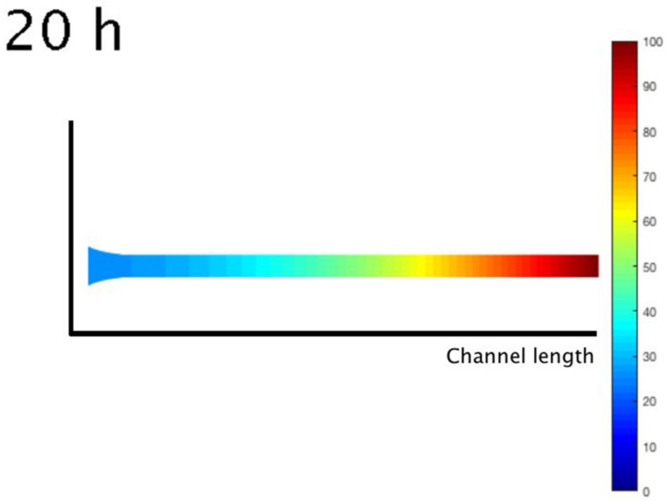


**Movie S2. FUCCI-MDA-MB-231 cells migration on 2D gelatin.**

Time lapse of two representative FUCCI-MDA-MB-231 cells in G1 phase (left, red nucleus) and S/G2 phase (right, green nucleus) of the cell cycle. Corresponding centroid tracks are shown in red and green. Time between frames is 10 minutes. Scale bar is 50 µm.

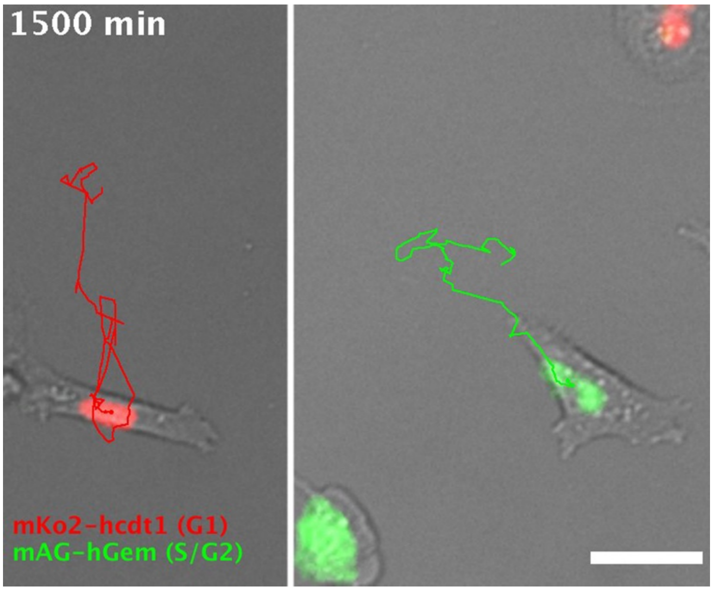

**Movie S3. FUCCI-MDA-MB-231 cells migration inside the microchannels.**

Time lapse of two representative FUCCI-MDA-MB-231 cells in G1 phase (bottom, red nucleus) and S/G2 phase (top, green nucleus) of the cell cycle migrating inside microchannels. Centroid tracking is shown in red and green. Time between frames is 10 minutes. Scale bar is 50 µm.

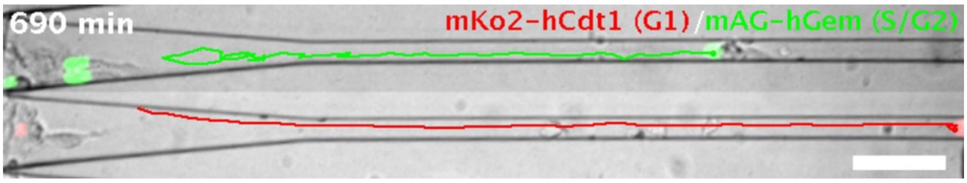


**Movie S4. Computational image segmentation and tracking performed by LEVER**

The movie is a merged view of mKO2-hcdt1 (red, nucleus marker of G1 phase), mAG-hGEM (green, nucleus marker of S/G2 phase) and phase channels. Left panel depicts a time lapse movie of FUCCI-MDA-MB-231 cells migrating inside vertically aligned fibers of 3D collagen. Right panel demonstrates results of cell segmentation via LEVER. Time between frames is 10 minutes.

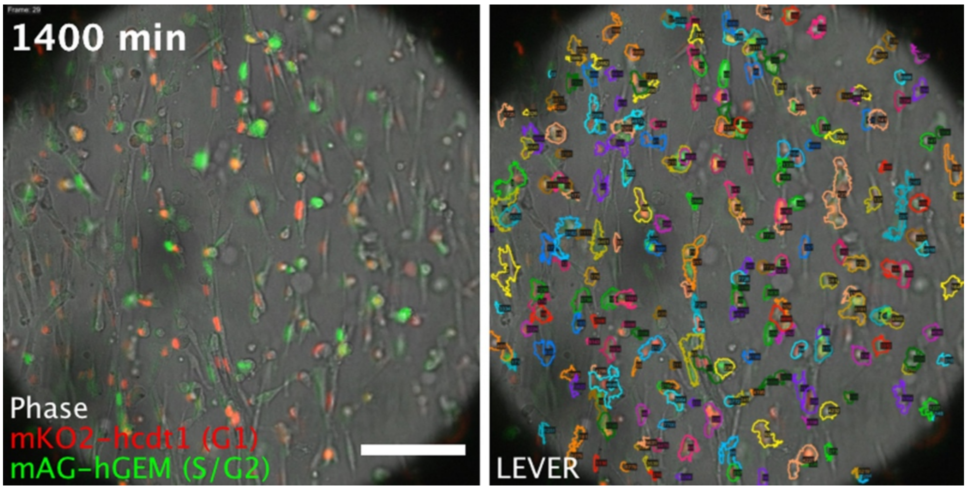


**Movie S5. FUCCI-MDA-MB-231 cells migration in 3D collagen with random fiber structure**

Time lapse of two representative FUCCI-MDA-MB-231 cells in G1 (red) and S/G2 (green) phase of the cell cycle migrating within the isotropic 3D collagen fibers. Corresponding centroid tracks are shown in red and green. Time between frame 10 minutes. Scale bar is 50 µm.

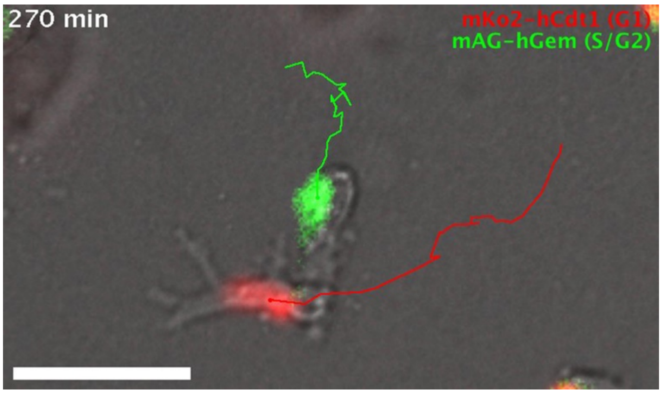

**Movie S6. FUCCI-MDA-MB-231 cells migration in 3D collagen with vertically aligned fiber structure**

Time lapse of two representative FUCCI-MDA-MB-231 cells in G1 (red, left) and S/G2 (green, middle) phase of the cell cycle migrating in vertically aligned 3D collagen fibers. Right panel is a merged view of red (mKO2-hCdt1), green (mAG-hGEM) and phase channels overlaid with corresponding centroid tracks. Time between frame 10 minutes. Scale bar is 50 µm.

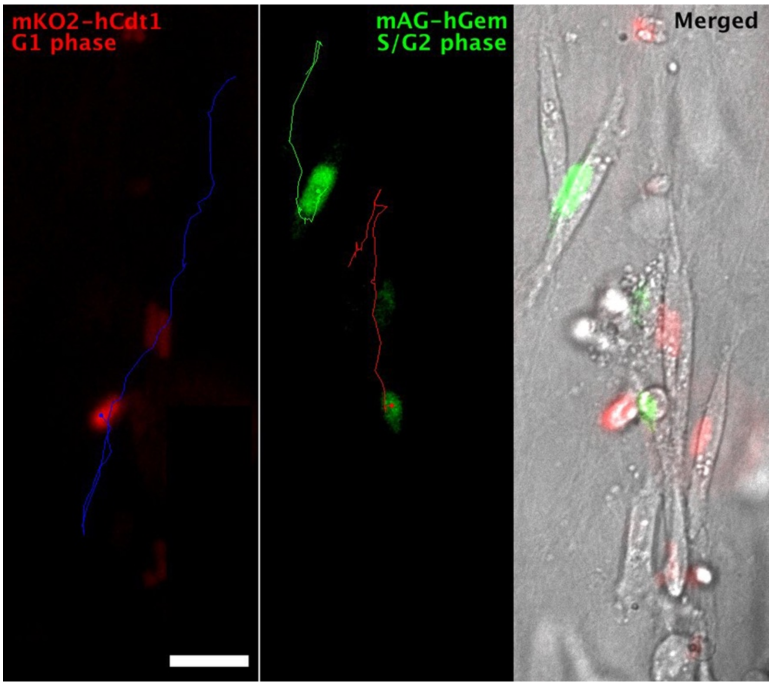
